# Lysyl oxidase inhibition drives the transdifferentiation from lung adenocarcinoma to squamous cell carcinoma in mice

**DOI:** 10.1101/314393

**Authors:** Shun Yao, Xiangkun Han, Xinyuan Tong, Fuming Li, Zhen Qin, Hsin-Yi Huang, Ji Hongbin

**Affiliations:** State Key Laboratory of Cell Biology; Innovation Center for Cell Signaling Network; CAS Center for Excellence in Molecular Cell Science, Shanghai Institute of Biochemistry and Cell Biology, Chinese Academy of Sciences, Shanghai, 200031, China; University of Chinese Academy of Sciences, Beijing, 100049, China; School of Life Science and Technology, Shanghai Tech University, Shanghai, 200120, China

**Keywords:** Lung adenocarcinoma, Squamous cell carcinoma, Lysyl oxidase, Transdifferentiation, LKB1

## Abstract

*LKB1* is frequently mutated in human non-small cell lung cancer (NSCLC) and *Lkb1* deletion in mice triggered the lung adenocarcinoma (ADC) to squamous cell carcinoma (SCC) transdifferentiation (AST) through lysyl oxidase (LOX)-dependent extracellular matrix remodeling. Here we show that pharmacological inhibition of lysyl oxidase in *Kras^G12D^/Trp53^L/L^* mouse model, which is known to produce lung ADC only, triggers the ADC-to-SCC transdifferentiation independent of *LKB1* status. Treatments of two different inhibitors of lysyl oxidase decrease collagen deposition and promote redox accumulation, and eventually trigger the AST. Importantly, these transited SCC show strong resistance to lysyl oxidase inhibition in stark contrast to ADC. Collectively, these findings establish a new AST mouse model independent of *LKB1* inactivation status.

## Introduction

Lung adenocarcinoma (ADC) and squamous cell carcinoma (SCC) are two distinct subtypes of non-small cell lung cancer (NSCLC). Interestingly, the mixed pathologies including both lung ADC and SCC in single lesions, clinically termed as adeno-squamous carcinoma (Ad-SCC), is detected in approximately 1-10% of lung cancer [1]. This indicates a potential phenotypic transition between lung adenomatous and squamous components in this pathologically mixed lung cancer.

Loss-of-function mutations of *LKB1*, also named Serine/Threonine Kinase 11 (*STK11*) have been frequently observed in human lung ADC, SCC as well as Ad-SCC specimens [2]. Using genetically engineered mouse models (GEMMs), we have previously found that *LKB1* deficiency is sufficient to drive lung ADC-to-SCC transdifferentiation (AST) [3, 4]. Interestingly, clinical specimen analyses show that up to 30% of human lung Ad-SCC indeed harbor *LKB1* genetic alterations and these tumors display strong plasticity, featured with frequent mixed pathologies [2, 4]. However, these data also indicate that the majority (~70%) of human lung Ad-SCC might have phenotypic transition routes independent of *LKB1* inactivation status.

Our previous study in *Kras^G12D^/Lkb1^L/L^* (*KL*) mouse model has demonstrated that lysyl oxidase (LOX) level dramatically decreases during the AST process, which is accompanied by down-regulation of collagen deposition and extracellular matrix (ECM) remodeling [3]. 3-Aminopropionitrile (BAPN) [5] and D-penicillamine (DPA) [6] are two well-documented chemicals that can inhibit LOX activity through different mechanisms: BAPN directly binds to the active site of LOX and irreversibly inhibits its enzymatic activity to affect the crosslinking of collagen molecules [7, 8] whereas DPA blocks LOX activity mainly through chelating its cofactor copper [9]. In *KL* mouse model, we find that either BAPN or DPA treatment significantly accelerates the AST process through decreased collagen deposition and ECM remodeling [3]. These data highlight the essential role of LOX activities as well as collagen deposition and ECM remodeling in lung AST process triggered by LKB1 loss in mouse model. These findings are further supported by clinical observation showing that a large portion of human lung SCC show decreased LOX level and collagen deposition than ADC [3].

The *Kras^G12D^/Trp53^L/L^* (*KP*) mouse model is widely accepted as closely recapitulating this subtype of human lung ADC [10]. In consideration of the important contribution of LOX inhibition in AST process in KL model, we want to test if BAPN or DPA treatment could promote similar phenotypic transition in *KP* mouse model, which is well-established to produce lung ADC only [7, 8]. Our data clearly show that LOX inhibition could trigger the squamous transdifferentiation of KP lung ADC independent of LKB1 status and thus establish a new AST mouse model.

## Results

We treated *KP* mice with BAPN or DPA daily via intraperitoneal injection post 4 weeks of Ad-Cre infection for 6-8 weeks and then harvested mouse lungs for pathological analyses (Fig. 1A). As expected, BAPN/DPA treatments resulted in dramatic reduction of collagen deposition in lung tumors (Fig. 1B). Moreover, we found that BAPN/DPA treatments significantly decreased lung tumor number as well as tumor burden (Fig. 1C) [11]. Lung tumors from control group exclusively displayed adenomatous pathology and expressed adenocarcinoma biomarkers such as TTF1 and SP-C (Fig. 1D). Intriguingly, we found that certain lung tumors from treatment groups displayed typical squamous pathology, which featured with the expression of squamous markers p63 and K14 (Fig. 1C-D). The squamous tumors in BAPN treatment group were negative for SPC and TTF1 staining (Fig. 1D). In contrast, the squamous tumors in DPA treatment group showed negative SPC staining but some positivity for TTF1 staining (Fig. 1D). This is consistent with our previous finding of the TTF1 and p63 double positive squamous tumors in *KL* model [4]. Since it’s well established that *KP* tumors are uniformly with adenomatous pathology [10], our data demonstrated that pharmacological LOX inhibition triggered the phenotypic transition from lung adenocarcinoma to squamous cell carcinoma in *KP* mouse model.

**Figure 1.**
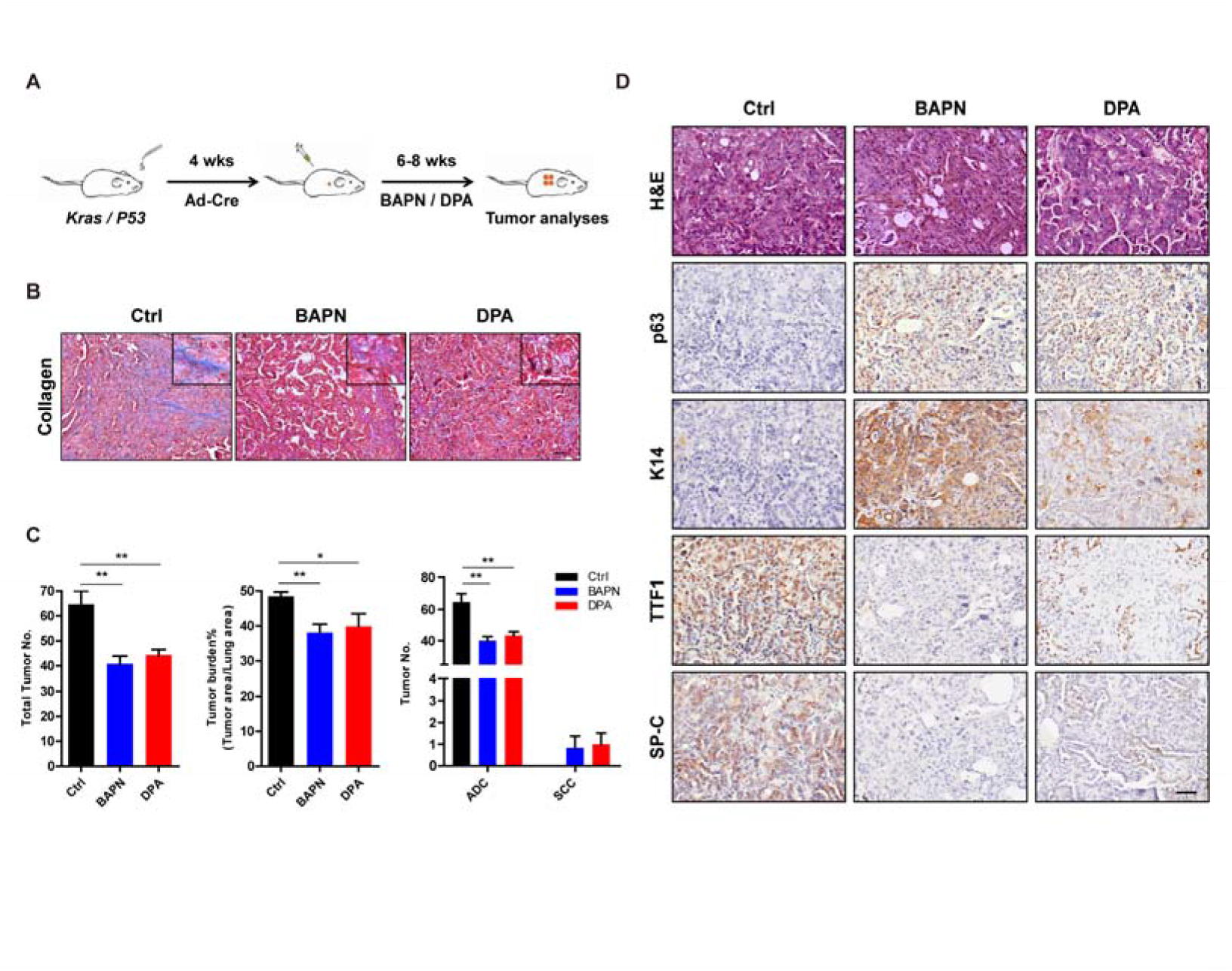
Pharmacological LOX inhibition promotes the ADC-to-SCC transdifferentiation in *KP* mouse model. **A**, Scheme for BAPN and DPA treatments in *KP* mouse model. **B**, Representative images of Masson’s Trichrome staining for collagen in *KP* mice treated with or without BAPN or DPA (six mice for each group). Scale bar, 50μm. **C**, Quantification of total tumor number (left), tumor burden (middle), ADC or SCC number (right) in *KP* mice treated with or without BAPN or DPA (six mice for each group). Data were shown as mean ± SEM. One-way ANOVA test followed by t test (left and middle), two-way ANOVA test followed by t test (right). ***P<0.001, **P<0.01, *P<0.05. **D**, Representative images showing H&E staining and IHC staining for p63, K14, TTF1 and SP-C in *KP* mice treated with or without BAPN or DPA. In BAPN/DPA treatment groups, the images only show the transdifferentiated SCC. Scale bar, 50μm.

*Lkb1* deficiency is the only well-established factor in driving AST in mice [3, 4]. We then analyzed whether LKB1 and its downstream signaling such as AMPK phosphorylation, which is known important for AST in *KL* model, were down-regulated during the AST process in *KP* model. Real time PCR and western blot data showed that both LKB1 mRNA and protein levels did not significantly change between BAPN/DPA treatment group and control group (Fig. 2A-B). This was further supported by LKB1 immunostaining in mouse lung tumors (Fig. 2C). Moreover, immunostaining analyses showed that AMPK phosphorylation, indicative of LKB1 activation status, remained comparable between control and treatment groups (Fig. 2C). Thus, these data supported that, in *KP* mouse model, the AST process triggered by LOX inhibition was independent of *LKB1* inactivation status.

**Figure 2.**
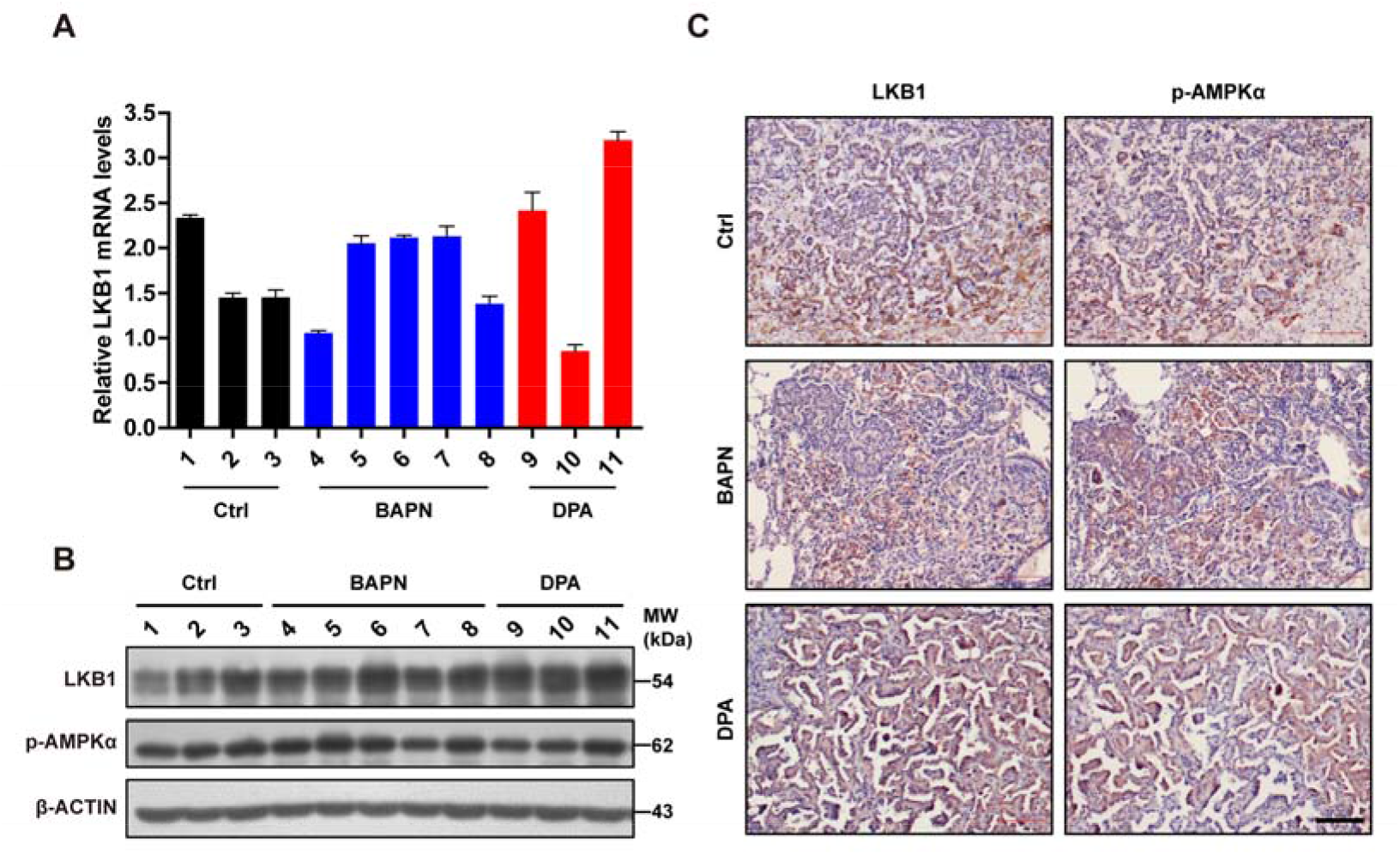
LKB1 and p-AMPKα levels in *KP* mice treated with LOX inhibitors. **A**, Relative mRNA levels of LKB1 in lung tumors of *KP* mice treated with or without BAPN or DPA. Data were shown as mean ± SEM. **B**, Western blotting analysis of LKB1 and p-AMPKα in lung tumors of *KP* mice treated with or without BAPN or DPA.-ACTIN served as internal control. **C**, Representative images for LKB1, p-AMPKα IHC staining in *KP* mice treated with or without BAPN or DPA. Scale bar, 100μm.

We further found that BAPN/DPA treatments in *KP* model promoted cell apoptosis and inhibited cell proliferation in lung ADC (Fig. 3A), consistent with our previous findings in *KL* mouse model [3]. Interestingly, we found the transited SCC showed significantly decreased apoptosis rate and almost no growth arrest in response to BAPN/DPA treatments (Fig. 3B-C). This pathology-specific drug response pattern indicated that the AST process might be an intrinsic strategy for lung cancer cells to adapt to ECM remodeling and escape treatments. ECM deprivation is a major cause of oxidative stress that contributes to the accumulation of reactive oxygen species (ROS) [12]. We then checked the level of 8-hydroxydeoxyguanosine (8-oxo-dGuo), the marker indicative of DNA oxidative modification (Fig. 4A). We found that the 8-oxo-dGuo level was significantly lower in SCC than ADC (Fig. 4A-B). These results demonstrated that LOX inhibition triggered the AST process might confer lung tumors with strong survival capability under environmental stress and promote the resistance.

**Figure 3.**
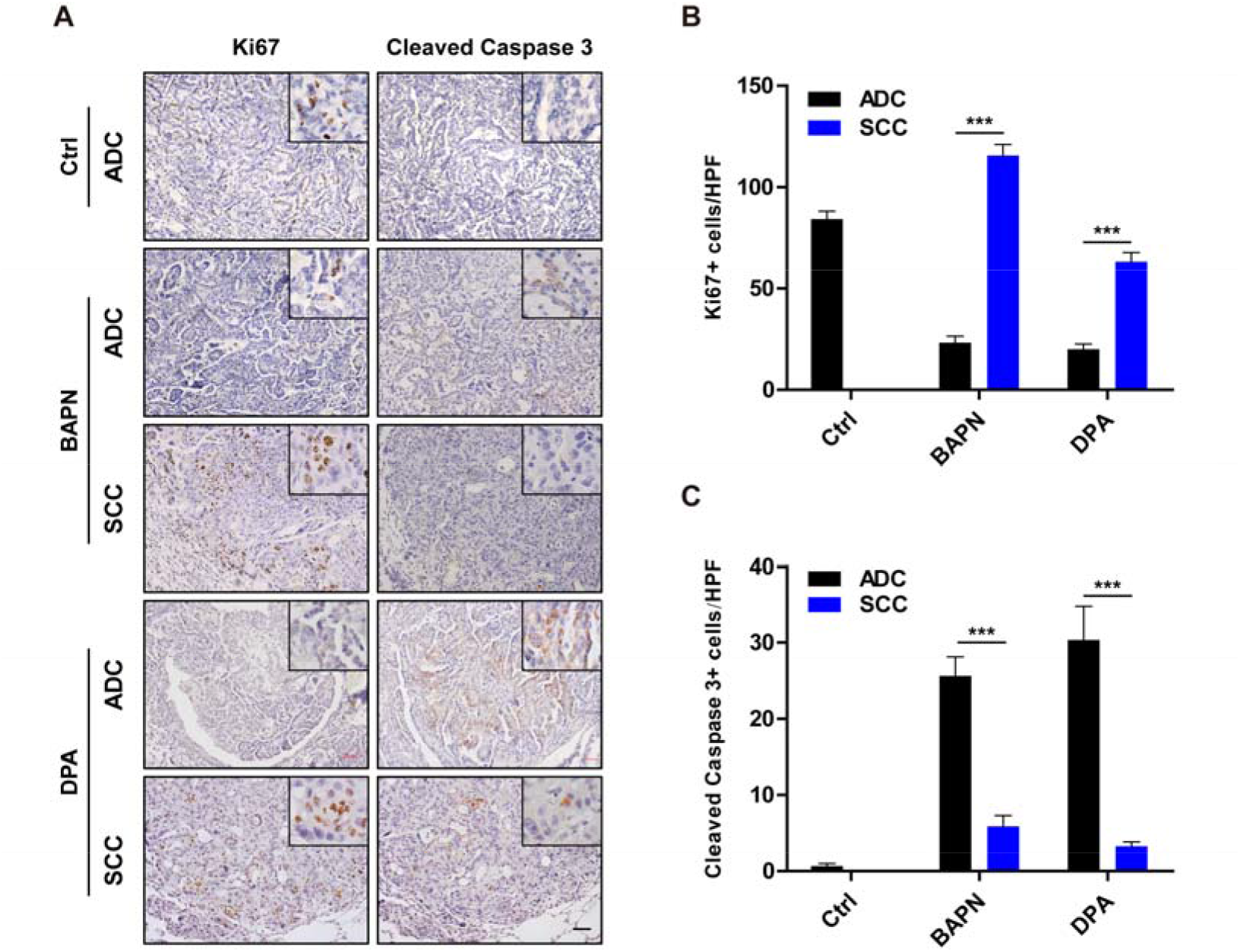
Ki67 and cleaved Caspase 3 immunostaining in *KP* mice treated with LOX inhibitors. **A**, Representative images for Ki67 and cleaved Caspase 3 immunostaining in *KP* mice treated with or without BAPN or DPA (six mice for each group). Scale bar, 50 m. **B-C**, Statistical analyses of the Ki67+ cells (B) and cleaved Caspase 3+ cells (C) per high-power field (HPF) in *KP* mice treated with or without BAPN or DPA. Data were shown as mean ± SEM. t-test, ***P<0.001.

**Figure 4.**
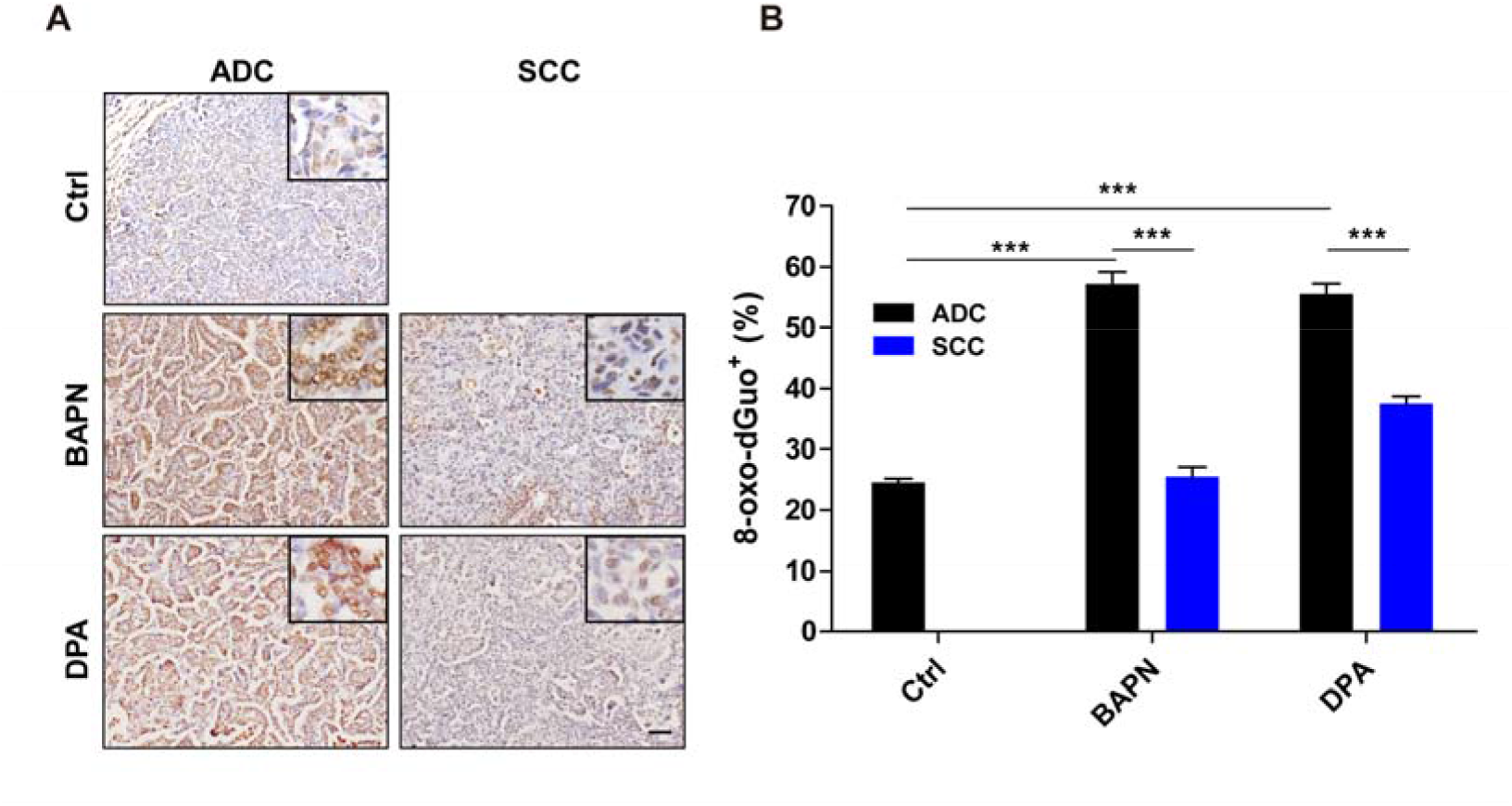
Immunostaining of 8-oxo-dGuo in *KP* mice treated with LOX inhibitors. **A**, Representative images for 8-oxo-dGuo immunostaining for ADC and SCC in *KP* mice treated with or without BAPN or DPA (six mice for each group). Scale bar, 50 m. **B**, Quantification of 8-oxo-dGuo for ADC and SCC in *KP* mice treated with or without BAPN or DPA. Data were shown as mean ± SEM. Two-way ANOVA test followed by t-test. ***P<0.001, **P<0.01, *P<0.05.

## Discussion

Genetically engineered mouse models, closely recapitulating human lung cancers, have been proven very helpful in understanding tumor initiation, progression as well as drug responses. Most previous GEMM have recapitulated lung cancer with adenomatous pathology, such as LSL-Kras^G12D^ [13], EGFR L858R [14]. Recent studies have also established several lung squamous cell carcinoma mouse models, e.g., *Lkb1^f1/f1^;Pten^f1/f1^* mice [15], *Sox2* overexpression in *Pten^f1/f1^;Cdkn2ab^−/−^* mice [16], transformation of airway basal cell from *Trp53^f1/f1^/Keap1^f1/f1^* mice [17]. In contrast, the *Kras^G12D^/Lkb1^L/L^* mouse model represents as the only well-established Ad-SCC model, which displays multiple lung cancer pathologies including ADC, SCC and mixed Ad-SCC [2]. Lung ADC from *KL* model can progressively transit to SCC via mixed Ad-SCC pathology potentially through LOX down-regulation, which results in decreased collagen deposition and ECM remodeling [3, 4].

We here find that the enzymatic inhibition of LOX could promote the ADC to SCC transdifferentiation in *KP* mice, which has wild type Lkb1 alleles. We show that both lung ADC and SCC from *KP* model show no significant change of LKB1 gene expression as well as down-stream AMPK phosphorylation. This supports that such AST triggered by LOX inhibition is potentially independent of LKB1 inactivation status. It’s worth noting that the SCC transition frequency in this *KP* model is quite low. We reason this could be due to the duration of LOX inhibitor treatment and a longer BAPN/DPA treatment could further promote the squamous transdifferentiation. Our data show that LOX inhibition results in decreased collagen deposition and ECM remodeling. ECM depletion is known to promote the accumulation of reactive oxygen species [12] and important for triggering AST via currently unknown mechanism [4]. Consistently, we find that treatments with LOX inhibitors significantly increase 8-oxo-dGuo, a DNA oxidation modification marker, indicating the excessive ROS accumulation in KP ADC. This might explain the squamous transdifferentiation of lung ADC in *KP* model.

In *KL* model, the transited SCC become more resistant to the treatment of small compounds such as phenformin [4]. Similar observations are also detected in *KP* model when treated with LOX inhibitors. In contrast to ADC, the transited SCC in *KP* model also show much higher proliferation rate and lower apoptosis rate. Multiple studies including our own work have convincingly demonstrated that LOX promotes cancer metastases through promoting collagen deposition and targeting LOX might serve as a novel therapeutic strategy [18–21]. Consistent with this, we have observed that lung ADC from *KP* model show significant tumor regression in response to BAPN/DPA treatments. However, the SCC seems resistant to LOX inhibition, indicative of the pathological-specific therapeutic effects of LOX inhibitors. Indeed, multiple recent clinical studies have supported that histological transformation from lung ADC to SCC might contribute to therapeutic resistance [22]. Besides squamous transdifferentiation, phenotypic transition to neuroendocrine cancer has also been clinically implicated in drug resistance prostate cancer [23]. These data together indicate that pathological transition might emerge as an important factor in determining drug response.

## Materials and Methods

### Mouse Colony, Mouse Treatment, and Tumor Analyses

*Kras^G12D^/Trp53^L/L^* (*Kras/Trp53*) mice were originally generously provided by Drs. Tyler Jacks and Kwok-Kin Wong. Mouse care and treatment was approved by the Animal Care and Use Committee at the Institute of Biochemistry and Cell Biology, Shanghai Institutes for Biological Sciences, Chinese Academy of Sciences. Mice were treated at 6–8 weeks of age via nasal inhalation of adenovirus carrying Cre recombinase (2×10^6^ PFU).

For pharmacological treatments in Kras/Trp53 mice, either BAPN (100 mg/kg), DPA (150 mg/kg) or saline were given to mice at 4 weeks post Ad-Cre treatment via intraperitoneal injection daily as previously described [3]. LOX inhibitors: 3-Aminopropionitrile fumarate salt (A3134-10G, Sigma-Aldrich), DPA (P4875-5G, Sigma-Aldrich). The treatment period lasted for 6-8 weeks. Mice were then sacrificed and the whole lungs were fixed with 4% paraformaldehyde for the preparation of routine paraffin section. Hematoxylin and eosin (H&E) staining were performed as described previously [3]. Tumor number was counted under microscope and tumor size was analyzed using Image J software [14].

### Immunohistochemistry staining and antibodies

Mouse lungs were inflated with 1ml 4% paraformaldehyde, fixed overnight and dehydrated in ethanol, embedded in paraffin, sectioned at 5μm followed by staining with haematoxylin and eosin. Immunohistochemistry was performed as previously described [2, 21]. The following antibodies were used: SP-C (AB3786, Chemicon, 1:2000), p63 (SC-8431, Santa Cruz, this antibody specifically recognizes DNp63; 1:200), LOX (L4669, Sigma, 1:800), TTF1 (5883-1, Epitomics, 1:500), K14 (PRB-155P, Covance, 1:1000), Ki67 (NCL-Ki67p, Leica, 1:500), cleaved Caspase 3 (9664, Cell Signaling, 1:500), 8-oxo-dGuo (ab48508, Abcam, 1:300), pho-AMPK Thr172 (2535, Cell Signaling, 1:100), LKB1 (13031, Cell Signaling, 1:100).

### Masson’s Trichrome Collagen staining

Masson’s trichrome stain was performed to assess the collagen deposition. The tissue specimens were fixed in 4% paraformaldehyde, routinely paraffin embedded, and sliced into 5μm thick sections. The Masson’s trichrome staining procedures are as follows: sections are deparaffinized to water. Then stain the tissues with Weigert’s working hematoxylin for 10 minutes and blue in running tap water. Biebrich scarlet stains for 5 minutes and rinse in distilled water. Phosphotungstic/phosphomolybdic acid treats for 10 minutes, and transfer the sections directly into Aniline blue for 5 minutes. Rinse the sections in distilled water and wash with 1% Acetic acid for 1 minute. Rinse with distilled water, dehydrate, and cover slip. The results should be: nuclei—black; cytoplasm, muscle, erythrocytes—red; collagen—blue.

### Western Blotting

The western blotting was performed as previously described [3, 21]. Briefly, whole cell lysates were prepared in lysis buffer. Equal amounts of protein were resolved by electrophoresis on gradient gels (Bio-Rad). The proteins were electro-transferred to PVDF membrane (Pall). Primary antibodies were incubated at 4°C overnight and the HRP-conjugated secondary antibodies were incubated at room temperature for 1 hr. The following antibodies were used: pho-AMPK Thr172 (2535, Cell Signaling, 1:1000), LKB1 (13031, Cell Signaling, 1:500), β-ACTIN (A2228, Sigma, 1:5000).

### Reverse Transcription and Quantitative PCR Analysis

Equal amounts of total RNA were reverse transcribed into first-strand cDNA using Revert Aid™ First Strand cDNA Synthesis Kit (Fermentas). The cDNAs were used for real time PCR on a 7500 Fast Real-Time PCR System (Applied Biosystems) using SYBR-Green Master PCR mix (Roche). β-actin served as internal control. The primers for Q-PCR were summarized in the following table.

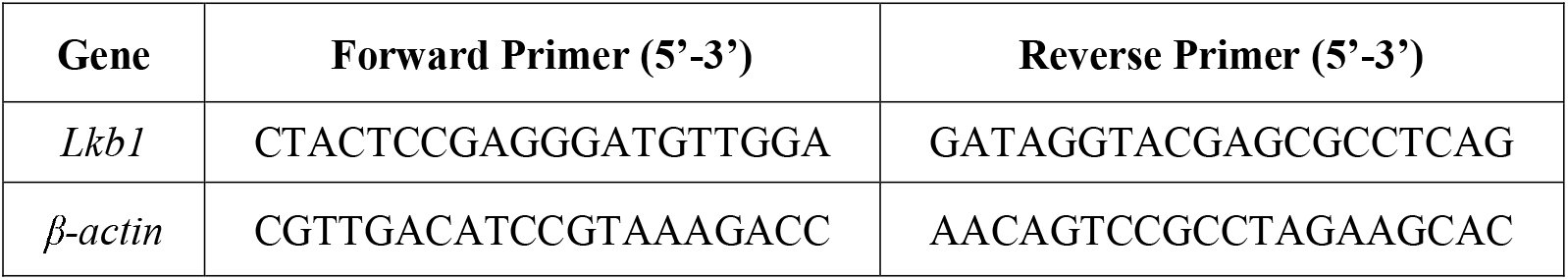

### Statistical analysis

Differences were compared using ANOVA and Student’s t-test. P-value<0.05 was considered statistically significant. All analyses were performed with Graph Pad Prism 5 software.

## Acknowledgments

The authors thank Dr. Tyler Jacks (Cambridge, MA) and Dr. Kwok-Kin Wong (Boston, MA) for kindly providing the *Kras/Trp53* mice.

## Authors’ Contributions

**Conception and design:** S. Yao, X.K. Han, H. Ji

**Development of methodology:** S. Yao, X.K. Han, F.M. Li, H. Ji

**Acquisition of data (provided animals, acquired and managed patients, provided facilities, etc.):** S. Yao

**Analysis and interpretation of data (e.g., statistical analysis, biostatistics, computational analysis):** S. Yao, X.K. Han

**Writing, review, and/or revision of the manuscript:** S. Yao, X.K. Han, H.Y. Huang, H. Ji

**Administrative, technical, or material support (i.e., reporting or organizing data, constructing databases):** S. Yao, X.K. Han, H.Y. Huang, X.Y. Tong, Z. Qin, H. Ji

**Study supervision:** H. Ji

## Disclosure of Potential Conflicts of Interest

The authors declare no potential conflicts of interest.

## Notes

**Grant Support** This work was supported by the National Basic Research Program of China (2017YFA0505500), the Strategic Priority Research Program of the Chinese Academy of Sciences, Grant No. XDB19000000, the National Natural Science Foundation of China 81430066, 91731314, 31621003), Science and Technology Commission of Shanghai Municipality (15XD1504000) and China Postdoctoral Science Foundation (2015M581673).

